# Bias in genome-wide association test statistics due to omitted interactions

**DOI:** 10.1101/2025.11.21.689603

**Authors:** Burak Yelmen, Merve Nur Güler, Estonian Biobank Research Team, Tõnu Kollo, Märt Möls, Guillaume Charpiat, Flora Jay

## Abstract

Over the past two decades, genome-wide association studies (GWAS) enabled the discovery of thousands of variants associated with many complex human traits. However, conventional GWAS are still widely performed with linear models with the assumption that the genetic effects are predominantly additive. In this work, we investigate the test statistic behavior when linear models are used to obtain significant genotype-phenotype associations without accounting for epistasis. We first algebraically derive mean and variance shift in the null statistic due to the omitted interaction term, and define the boundary between conservative (i.e., deflated statistic tail) and anti-conservative (i.e., inflated statistic tail) regimes for the common GWAS significance threshold. We then perform phenotype simulation analyses using the Estonian Biobank genotypes and validate the mathematical model. We demonstrate that the anti-conservative regime is plausible under realistic parameter settings and models omitting interaction terms can produce spurious significance. Our findings suggest caution when interpreting statistically significant signals reported in the literature based on linear models, especially for large-scale GWAS.

## Introduction

Biological systems consist of multiple interrelated components, which makes them difficult to model faithfully using linear functions. It is reasonable to assume that the relationship between genetics and complex traits exhibits similar complexity. Indeed, multiple studies have proposed epistasis and gene-environment interactions as fundamental causal mechanisms of phenotypic variance [1–3]. Genome-wide association studies (GWAS), despite allowing the discovery of thousands of genotype-phenotype links with functional importance, are still widely performed with linear models after almost two decades [4]. This prevailing methodological trend might be attributed to multiple factors such as the ease of interpretability, attainable computational requirements, tradition and studies suggesting that the genetic component of phenotypic variance can mainly be explained by additive models [5–7].

This assumption of mainly additive effects, however, may hinder the efforts to understand the causal genetic mechanisms since the true associations might be nonlinear, despite being captured without nonlinear modeling [8]. Even if we accept the mainly additive model for association discovery, recent methods point to large variability of non-additive variance (i.e., the left-over variance after accounting for additive variance) among phenotypes [9]. In this light, various works have either tried to approach GWAS with nonlinear models such as neural networks [10, 11] or explicitly infer/integrate epistasis [12–14].

There are some hints in the literature on the consequences of omitting interaction terms in GWAS. Omitted variable bias, which is the bias in model-estimated coefficients due to the exclusion of variables relevant to the response variable, is a well known statistical concept [15] and it has been used to demonstrate that GWAS summary statistics might tag non-additive variance whenever there is nonzero correlation between genotypes and the interaction terms [9]. Furthermore, some evidence was presented in a recent study for “synthetic associations” (i.e., associations due to combined effects of multiple other variants [16, 17]) due to epistasis [18]. Interestingly and perhaps alarmingly, it was suggested that 3-5% of GWAS Catalog peaks might be synthetic, but the findings were obtained from indirect evidence based on unusual linkage disequilibrium (LD) patterns and did not necessarily point to spurious associations due to epistasis. Here we use “spurious” to refer to associations arising purely due to epistasis which does not include the target SNP. To the best of our knowledge, the effects of omitting interaction terms in GWAS have not been modeled to understand whether this omission can result in spuriously significant associations under realistic settings.

In this work, we derive a mathematical model to assess the effects of omitted interaction terms on GWAS test statistics. We initially define a true data generating process (DGP) for a phenotype with a fixed interaction component. Our mathematical setup is that a standard linear mixed model (LMM) is fitted for this phenotype to estimate the target SNP coefficient *α*. We preprocess all vectors from the true DGP and the fitted model to whiten and remove covariate effects. Based on the equality of the phenotype value for both models after preprocessing, we derive ordinary least squares (OLS) estimates for the expected value of the estimated 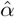 and its variance, under the null hypothesis *α* = 0. These allow us to derive the mean and variance of the true null statistic as opposed to the nominal null 𝒩 (0, 1). We then define the *R*(*x*) conservativeness ratio of true to nominal p-value to assess anti-conservative and conservative regimes for a given critical value *x*. To ensure that both the SNP and interaction effects are clearly defined, we introduce an extended null concept, strict no-path null, in which not only the target SNP coefficient is 0, but also the target SNP cannot be part of the interaction features. Intuitively, this makes sure that any identified bias of the test statistics is purely due to the realized interaction component and its correlation with the target SNP. Using this strict null concept, we simulate two sets of 10,000 phenotypes based on the Estonian Biobank genotype dataset [19] with varying non-additive variance fraction estimates from the literature, use REGENIE [20] to obtain summary statistics for target SNPs and confirm the mathematical model. Both the mathematical model and the simulation results demonstrate that even under minimalistic non-additive variance estimates and low correlation between the target SNP and the interaction component, anti-conservative regime is plausible and multiple non-associated SNPs are identified as significant due to the omitted interaction term. The key interpretation of our findings is that widely used linear models which do not compensate for epistatic effects can be prone to identifying spurious associations, especially when the sample size is large.

## Materials and Methods

### True data-generating process (DGP)

For individuals *i* = 1, …, *n*, let *y* ∈ ℝ^*n*^ denote the quantitative trait, *X* ∈ ℝ^*n*×*q*^ the observed covariates (e.g., age, sex, principal components etc.) and *g* ∈ ℝ^*n*^ the target SNP. The true DGP is

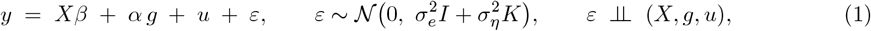

where *u* ∈ ℝ^*n*^ is the realized (fixed) mean contribution of possibly high-dimensional interaction terms (i.e., *u* = *Zβ*_*u*_ for some matrix *Z* ∈ ℝ^*n*×*m*^ whose columns encode SNP–SNP interaction terms) and *K* is the genetic relatedness matrix (GRM). We treat (*X, g, u*) as fixed and *ε* (containing environmental and additive polygenic effects) as stochastic.

### Fitted (misspecified) model

We fit the standard linear mixed model (LMM)

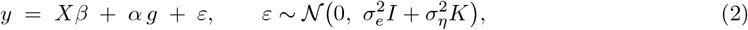

and examine the test *H*_0_ : *α* = 0 performed under (2) when the true DGP is (1).

### Preprocessing

Define a single linear operator *T* that whitens and removes covariates:

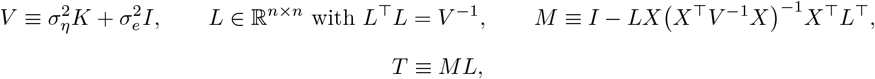

where *L* is the whitening matrix and *M* is the orthogonal projector, in the whitened space, that removes the subspace containing the covariate component *X*, that is, *TX* = 0. Note that *M* ^2^ = *M* . We assume *V* is (ideally) known/estimated correctly in a generalized least squares (GLS) framework and apply *T* to every vector in (1) and (2). From now on, we overwrite the symbols

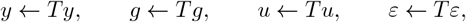

and work entirely with these transformed variables. In this space, the noise is spherical on the working subspace: *ε* := *T* 𝒩 (0, *V*) = *M* 𝒩 (0, *I*) and consequently Cov_*ε*_(*ε*) = *M*^*T*^ *M* = *M* . The fitted (misspecified) model (Eq. 2) thus becomes:

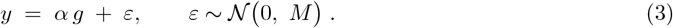

Moreover, this noise is almost unit-normed on average:

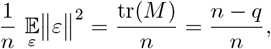

which is close to 1 for large *n*, as *q* ≪ *n* in typical set-ups. Note that under *H*_0_ : *α* = 0, the (true) working model reduces to

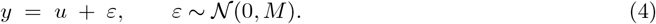

### Moments

Let us consider the empirical variance of *g*_*i*_ and *u*_*i*_ with respect to individuals *i*:

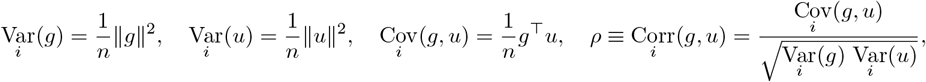

supposing centered variables. All moments are computed in the preprocessed space.

### Expected value of the coefficient

The no-intercept OLS estimate of *α* from the misspecified model (Eq. 3) is

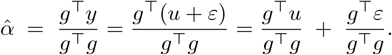

Hence, expected value of 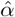 over *ε* is

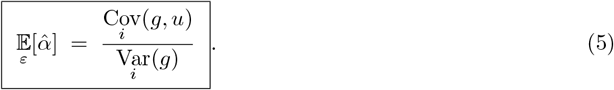

### Residual variance

The OLS residuals between the observed *y* (after preprocessing) and their explained component 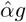 from the linear regression with the target SNP *g* are:

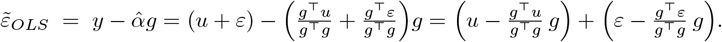

The two summands are uncorrelated on average over *ε*, yet spuriously slightly empirically correlated. Thus the residual empirical variance is the sum of their variances, plus a small covariance that vanishes with sample size:

1. Contribution from *u*:

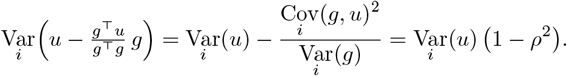
2. Contribution from *ε*: the empirical variance of the second term is

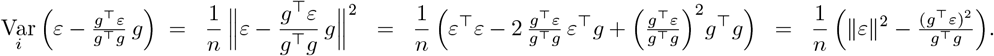

As 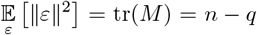, and 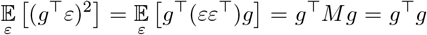 (since *g* is obtained by applying *T* = *ML* as preprocessing and *M* ^2^ = *M*), we obtain that on average, this variance over *ε* is:

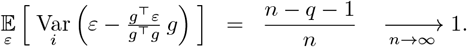

Moreover, when the number *n* of individuals is large, for any given realization of *ε*, the empirical variance (b) is likely to be very close to this average, because the variance w.r.t. *ε* vanishes (see proof in Appendix A):

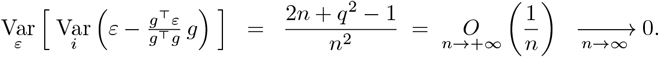
3. The spurious cross-contribution from *u* and *ε*:

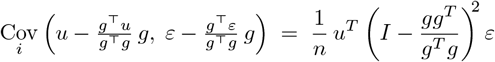

similarly vanishes with high sample size *n*. Indeed, its average over *ε* is 0, as *ε* is independent from *u* and *g*, and its variance w.r.t. *ε* also tends to be 0, being equal to 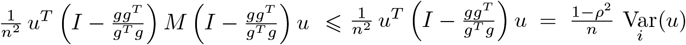, using the fact that the operators 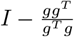 and *M* are projectors.

Hence the (asymptotic) total residual variance is

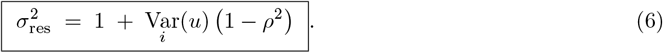

### Standard error

For no-intercept single-regressor OLS, asymptotically

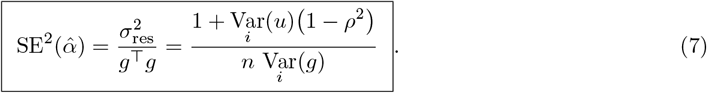

### The *t*-statistic

The null hypothesis *α* = 0 is tested with the *t*-statistic 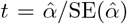. Under the working model, randomness arises only from *ε*, while *u* and *g* are fixed, so 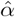 OLS estimator can be written as

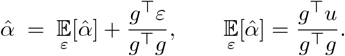

Therefore, conditionally on (*g, u*), the sampling variance of 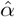 is

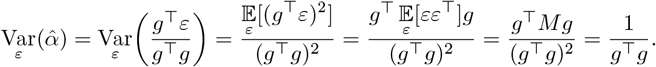

The OLS standard error, however, is based on the residual variance 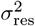 from (6) as derived in (7). Therefore, the variance of the *t*-statistic is asymptotically

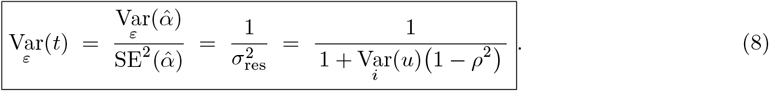

Consequently, the *t*-statistic is asymptotically 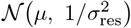 with

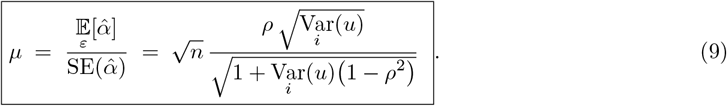

### Equivalent parameterization

Since the average coordinate-wise variance from *ε* is asymptotically 1 after preprocessing as in (6), we can define a parameter 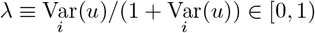, which is the proportion of variance explained by the interaction term. Then

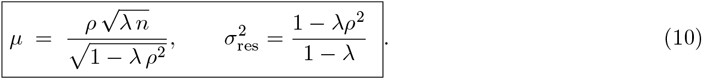

### Conservativeness

Let Φ denote the cumulative distribution function (CDF) of the standard normal distribution. Under the nominal null distribution 𝒩 (0, 1), the two-sided *p*-value for an observed statistic *x* is

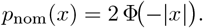

Under the true null distribution 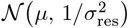, the actual two-sided tail probability is

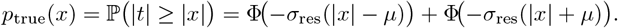

Then define the conservativeness measure

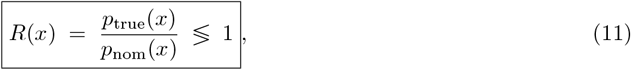

where *R*(*x*) > 1 indicates anti-conservativeness due to the shift in null mean and variance, and *R*(*x*) < 1 indicates conservativeness for a given statistic value *x*. In other words, for a critical value *x* (e.g., ± 5.45 corresponding to common GWAS significance threshold), *R*(*x*) > 1 means that observing *x* is more likely under the true null distribution, resulting in spurious significance.

### Strict no-path null

After preprocessing, decompose *u* as *u* = *u*_*¬g*_ + *u*_*g*_, where *u*_*¬g*_ aggregates interaction components not involving the target SNP and *u*_*g*_ those that do. Define the strict no-path null

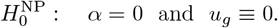

Under 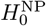 we have *u* = *u*_*¬g*_, so all expressions remain valid with *u* replaced by *u*_*¬g*_. Thus, any bias in the null *t*-statistic arises purely from alignment between *g* and omitted interaction signal that does not explicitly include *g* and does not involve any pathway in which *g* participates. Because *g* can be in LD with loci that contribute to *u*_*¬g*_, the correlation *ρ*_*¬g*_ can be nonzero even under this strict no-path null. Conversely, if *u*_*g*_ ≠ 0 (interactions that include *g*), then the shift includes a genuine path from *g* to *y* through nonlinear terms; in that case the single-variant test detects an association but its coefficient targets a best linear approximation rather than the additive parameter *α*. Therefore, in simulation analyses, we used this strict no-path null concept by excluding the target SNPs from the interaction features, to demonstrate only the effects of pure bias due to the omitted interaction term. It is important to emphasize here that this null concept was not chosen for realism but for controlled and well-defined evaluation of the bias. In reality, a target SNP might or might not be part of the interaction structure.

### Upper bound on *ρ*

We can provide plausible ranges for sample size *n* and the variance ratio *λ* from the literature, but *ρ* is difficult to estimate because the true interaction component for a given phenotype is unknowable. We can nevertheless obtain an upper bound without specifying interaction coefficients. Let *Z* ∈ ℝ ^*n*×*m*^ collect the allowed interaction features (e.g., two-way or three-way products), excluding any feature that explicitly uses the target SNP *g* (strict no-path). All columns are already in the preprocessed space (i.e., *Z* ← *TZ* and *g* ← *Tg* were applied upstream). Define the column space ℋ ≡ col(*Z*); under strict no-path we posit *u* ∈ ℋ. We seek the largest possible correlation between *g* and any interaction signal *u* consistent with the specification,

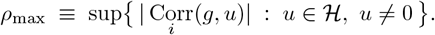

Let *P*_ℋ_ be the orthogonal projector onto ℋ (in the Euclidean inner product of the preprocessed space). Then

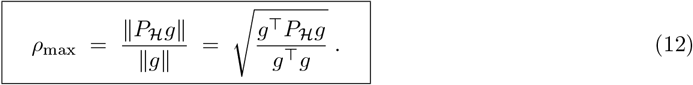

### Proof

Correlation is scale-free, so set ∥*u*∥ = 1. Then

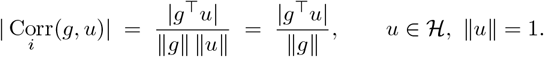

Decompose *g* as *g* = *P*_ℋ_*g* + (*I* − *P*_ℋ_)*g*, with *P*_*ℋ*_*g* ∈ ℋ and (*I* − *P*_ℋ_)*g* ⊥ ℋ. For any *u* ∈ ℋ,

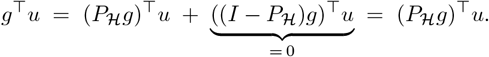

By Cauchy–Schwarz inequality, |(*P*_ℋ_*g*)^⊤^*u*| ≤ ∥*P*_ℋ_*g*∥ ∥*u*∥ = ∥*P*_ℋ_*g*∥, with equality for *u* = *±P*_ℋ_*g/*∥*P*_ℋ_*g*∥ whenever *P*_ℋ_*g* ≠ 0. Hence

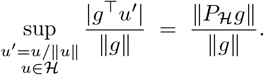

### Practical computation of *ρ*_*max*_

We compiled a dataset consisting of 210,145 samples with chromosomes 21 and 22 (total 8170 bi-allelic SNPs from GSA SNP array framework) from the Estonian Biobank [19] and mean-centered all SNPs. We used a list of unrelated individuals obtained with King v2.2.7 [21] to subsample to 100,144 individuals (removing up to second degree relatives). Using this filtered dataset, we can assume *V* has no structure. Therefore, we only residualized genotypes and *Z* to remove covariates (age, sex, first 5 principal components) and work approximately in the preprocessed space without the need for *V* estimation. We designed two different types of *Z* interaction matrices: one where target SNP and the interaction features are on the same chromosome and one where target SNP and interaction features are on different chromosomes. In both cases, the interaction feature matrix *Z* was generated randomly for 100 times with 100 two-way interaction terms, with the constraint that the target SNPs *g* are not part of these features (resulting in 383400 and 413500 total tested target SNPs, respectively, for the two designs over 100 iterations). To compute *ρ*_max_, we then solved the least-squares problem

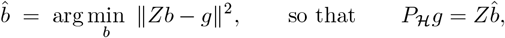

and obtained

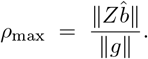

This allowed us to get *ρ*_max_ for a given interaction feature matrix *Z* without specifying the coefficients, hence without simulating phenotypes. Since *u* = *Zβ*_*u*_ and *β*_*u*_ ranges over an unbounded continuum, deriving *ρ*_max_ via *u* would otherwise require infinite simulations. Furthermore, we also assessed differences in the distribution of *ρ*_max_ estimates across *Z* feature matrices built from two-way to ten-way interaction terms (see Appendix B).

### Simulation analysis

Using the same genotype dataset of 210,145 samples, we simulated phenotypes as

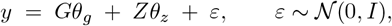

where *G* is the genotype matrix with 1000 SNPs on chromosome 22, *Z* is the two-way interaction matrix with 100 features (200 SNPs) formed with SNPs on chromosome 21, *θ*_*g*_ and *θ*_*z*_ are sampled from 𝒩 (0, 1) independently. To be able to imitate the previously defined variance fraction term *λ*, the terms in the simulation model were scaled so that the variance of the combined genetic and noise component would be 1. We varied the *λ* from 0.001 to 0.171, with 10,000 evenly spaced values, and for each value we drew a random genetic architecture as defined previously (i.e., total 10,000 simulated phenotypes). Furthermore, using these same steps, we obtained another set of 10,000 simulated phenotypes with the randomly subsampled sample size of 100,000. The upper limit for *λ* was obtained from Hivert et al. [7] estimation of upper limit for non-additive variance.

We then used a state-of-the-art LMM-based tool REGENIE v3.4 [20] to obtain test statistics over target SNPs, which were chosen to be on the same chromosome as the interaction SNPs while not being part of the interaction features (resulting in 3834 target SNPs for each simulated phenotype). This close proximity of target and interaction SNPs yielded multiple high-value realizations for the correlation term *ρ* with reasonable number of iterations. We provided age, sex and the first 5 principal components as covariates to REGENIE.

### Data availability

Code for reproducing the findings is available at https://github.com/genodeco/GWAS-Epistasis-Bias. Access to the Estonian Biobank dataset is subject to an application procedure, which can be found at https://genomics.ut.ee/en/content/estonian-biobank.

## Results

We first examined the distributions of *ρ*_*max*_ obtained only based on the interaction subspace and identified descriptive statistics such as the maximum *ρ*_*max*_ = 0.849 (when interaction SNPs and the target SNP are on the same chromosome) and *ρ*_*max*_ = 0.042 (when interaction SNPs and the target SNP are on different chromosomes) (Fig 1). Based on the mean, minimum and maximum values obtained from these distributions, we computed the conservativeness measure *R* from the expression (11), setting the *t*-statistic value to *x* = ± 5.45, which corresponds to the common GWAS significance p-value = 5e-8. For *ρ* values from 0.021 to 0.849, *R* was consistently observed in the anti-conservative regime (i.e., *R* > 1) over all *λ* and *n* values, and the regime where *p*_true_(*x* = ± 5.45) > 0.5 (i.e., an observation as extreme as |*x*| = 5.45 occurs at least 50% of the time, indicating spuriously significant hits at least ∼50% of the time) was achieved with sufficient *n* sample size (Fig 2). We further examined *n* versus *ρ* plots for *λ* = 0.055, *λ* = 0.119 and *λ* = 0.171 (estimated mean and maximum non-additive variance fractions from [7]), limiting *ρ* to *ρ*_*max*_ = 0.032 (Fig 3). *R* was again mostly observed in the anti-conservative regime. Furthermore, *p*_true_(*x* = ± 5.45) > 0.5 could be attained even with low *ρ* and *λ* values, and realistic sample sizes.

**Figure 1.**
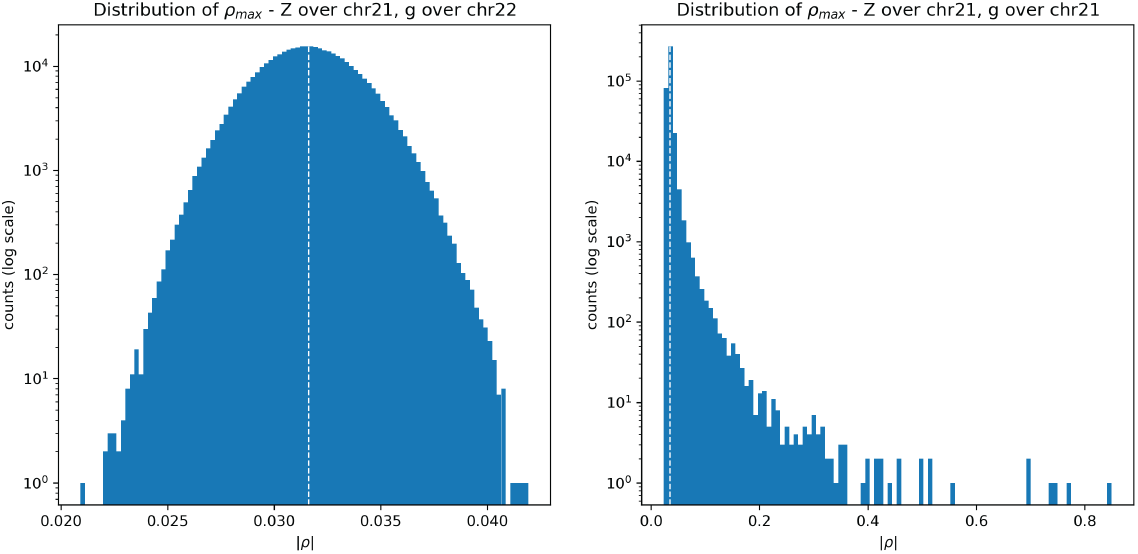
Distribution of *ρ*_*max*_ estimates obtained from the design where the interaction SNPs and target SNPs are on different chromosomes (left) and for the design where interaction SNPs and target SNPs are on the same chromosome (right). Vertical dashed lines indicate mean value. For the different chromosome design, min *ρ*_*max*_ = 0.021, mean *ρ*_*max*_ = 0.032, max *ρ*_*max*_ = 0.042. For the same chromosome design, min *ρ*_*max*_ = 0.024, mean *ρ*_*max*_ = 0.035, max *ρ*_*max*_ = 0.849.

**Figure 2.**
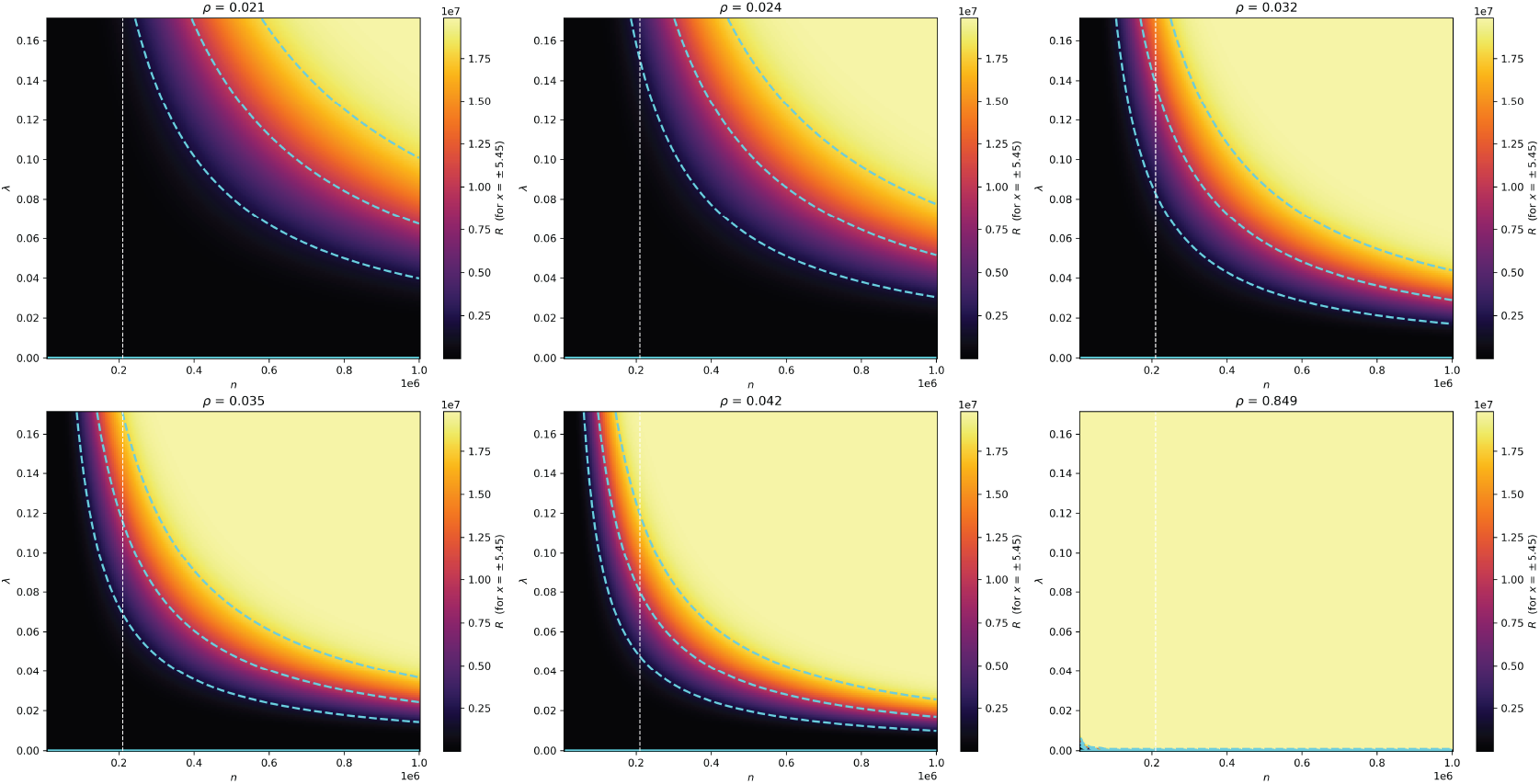
Heatmap plots of *R* = *p*_true_(*x*)*/p*_nom_(*x*) under the mathematical model for *x* = ± 5.45 with corresponding *n* (x-axis) and *λ* (y-axis) values, and different fixed *ρ* values. Cyan lines indicate *R* = 1 contour. Dashed cyan lines from center outwards indicate *p*_true_(*x*) = 0.1, 0.5 and 0.9, respectively. Vertical dashed line corresponds to sample size *n* = 210,145.

**Figure 3.**
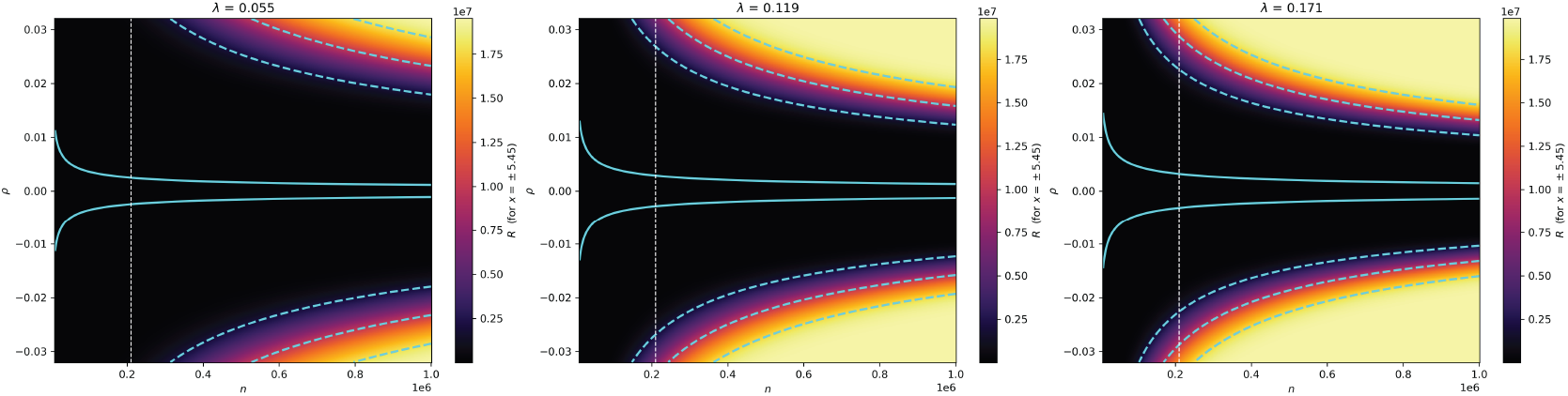
Heatmap plots of *R* = *p*_true_(*x*)*/p*_nom_(*x*) under the mathematical model for *x* = ± 5.45 with corresponding *n* (x-axis) and *ρ* (y-axis) values, and different fixed *λ* values. Cyan lines indicate *R* = 1 contour. Dashed cyan lines from center outwards indicate *p*_true_(*x*) = 0.1, 0.5 and 0.9, respectively. Vertical dashed line corresponds to sample size *n* = 210,145.

As the next step, we compared the results from REGENIE applied to the simulated phenotypes for *n* = 210,415 and *n* = 100,000 (by randomly subsampling the main dataset) with the mathematical model of conservativeness. There was high concordance between simulation results and the algebraic model for both *n* values (Fig 4). The expected spuriously significant hits were clearly observed in the simulations; mean |*z*| score was above 5.45 in the region corresponding to *p*_true_(*x* = ± 5.45) > 0.5 regime in the mathematical model (Fig 4a-b). Despite that the simulations did not produce some extreme value regions (such as high-*ρ* - low-*λ*) due to limited iterations, we were able to find clear evidence for *R* > 1 anti-conservative regime as we observed |*z*| > 5.45 for even extremely small *ρ* values (Fig 4c).

**Figure 4.**
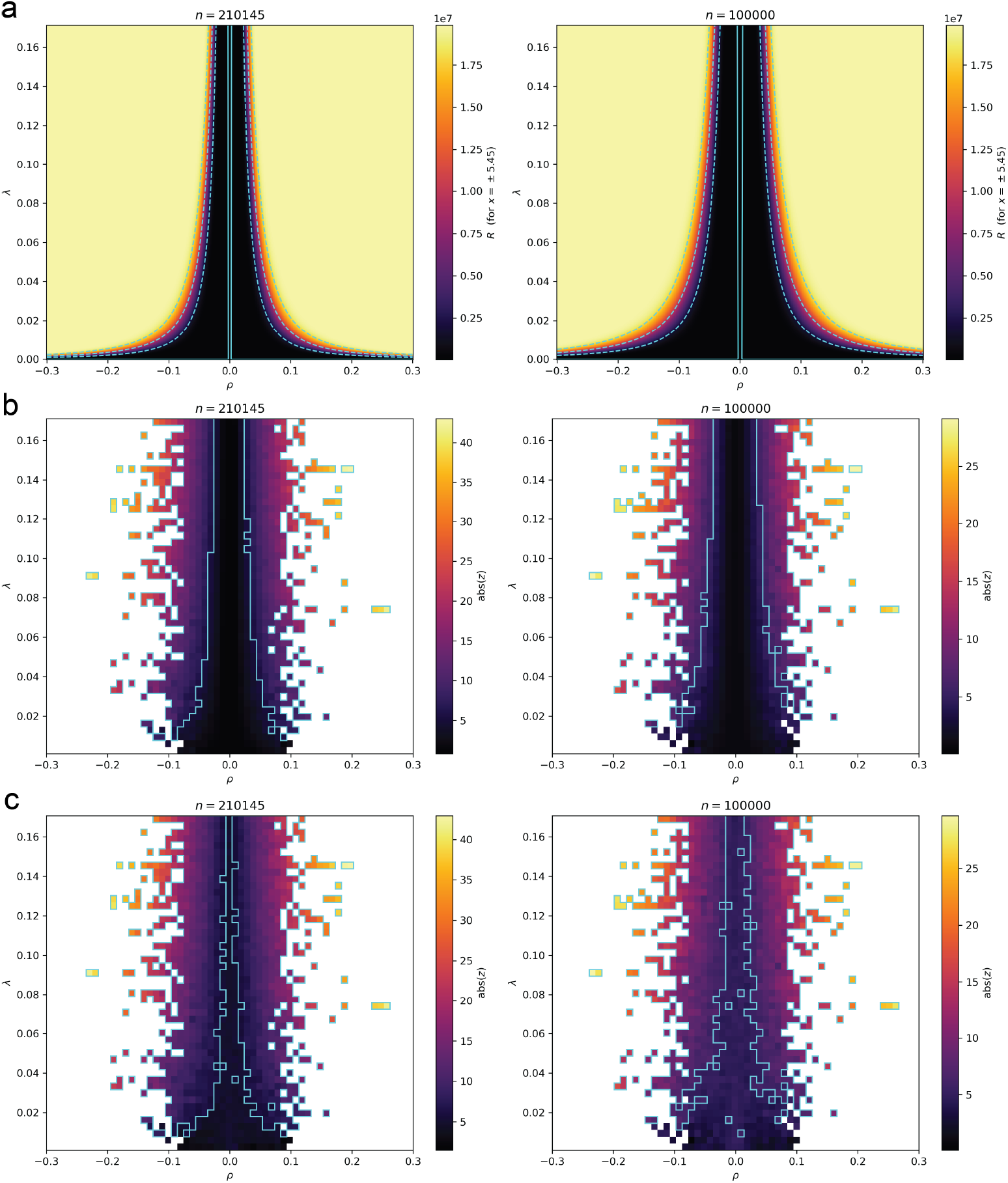
Comparison of mathematical model to simulated results for *n* = 100, 000 (right) and *n* = 210, 145 (left). **a.** *R* = *p*_true_(*x*)*/p*_nom_(*x*) ratio from the mathematical model with *x* = *±*5.45. Cyan lines indicate *R* = 1 contour. Dashed cyan lines from center outwards indicate *p*_true_(*x*) = 0.1, 0.5 and 0.9, respectively. **b**. Simulation-based heatmap plot of |*z*| scores obtained by mapping the values to a 50×50 grid and calculating the mean score for each square. White regions indicate squares where simulations did not have specific *λ* - *ρ* values (since *λ* could be set while simulating the phenotypes but *ρ* was only calculated depending on the realized interaction term). Cyan lines indicate |*z*| = 5.45 contour, corresponding to the common GWAS significance threshold. **c**. Simulation-based heatmap plot of |*z*| scores obtained by mapping the values to a 50×50 grid and calculating the maximum score for each square.

## Discussion

In this work, we derived a mathematical model characterizing the test statistic bias in GWAS if the true genetic architecture involves epistasis which is not taken into account in the fitted linear model. The inflationary bias at the extreme tail is mainly driven by the positive shift in the mean of the null statistic, whereas the downward shift in the variance drives deflation, yet the dominant tendency for the critical *t*-statistic value ± 5.45 (corresponding to the common GWAS significance p-value 5e-8) is inflation (hence, increased spurious significance) under a vast range of realistic parameter values (Fig 2, 3). We were able to define the mean and variance shift in terms of three terms: (i) *λ*, the variance fraction of the interaction component, (ii) *ρ*, the correlation between the target SNP and the realized interaction term, and (iii) *n*, the sample size. For the range of *λ*, we used the estimations from literature [7]. For the range of *ρ*, we first derived a maximum threshold algebraically without needing the interaction term coefficients. We also performed phenotype simulations with the full interaction architecture. As expected, even under the strict no-path null, absolute value of *ρ* can be high due to LD between the target SNP and the interaction SNPs. Interestingly, even when the target SNP and the interaction SNPs are on different chromosomes, *ρ* can reach sufficient non-zero values to produce spurious significance (Fig 1, 2). Another interesting observation was that the problem of spurious significance is positively correlated with the sample size. Average sample size for GWAS in 2022 was reported to be up to 140,000 [4]. It is now also not uncommon to utilize more than a million sample-sized datasets thanks to large biobanks [22]. As the sample size reaches a million, even minimal estimates for the interaction variance fraction *λ* (e.g., ∼0.03) and correlation *ρ* (e.g., ∼0.03) can result in *p*_true_(*x* = ± 5.45) ⪆ 0.5 regime where we expect approximately half of the statistically significant hits to be spurious (Fig 2).

One of the limitations of our model is the initial assumption of perfectly estimated *V* excluding *u*, which we mainly proposed to be able to equate true and misspecified model’s phenotype vectors. This is not realistic because the structured part of the estimated 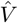 could potentially capture some variation from the interaction term. This effect heavily depends on how well the kinship matrix and polygenic variance estimations are affected by the unrealized true interaction structure. In other words, it depends on how much of the variance from the true interaction term is modeled under the non-genetic variance component of the estimated 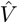. Modern LMMs generally estimate kinship matrix *K* and variance of the polygenic effect 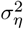 with the leave-one-chromosome-out (LOCO) method in which the chromosome of the target SNP is removed from the estimation to avoid proximal contamination. This means that our model’s limitation could especially be relevant if the target SNP and interaction SNPs are on different chromosomes, but the most severe bias where the target SNP and interaction SNPs are on the same chromosome most likely persists even when the *u* interaction space partially aligns with *K*. Indeed, we were able to confirm via LMM-based GWAS on simulated phenotypes that our mathematical model is consistent (Fig 4).

As a precautionary check for spurious significance without the omitted term, for each simulated phenotype, we also simulated a version for which the only difference is the subtraction of the realized interaction component from the final phenotype. We detected 4 and 2 spurious associations (i.e., p-val >5e-8) over 38,340,000 tested SNPs in 10,000 simulated phenotypes for *n* = 100, 000 and *n* = 210, 145 datasets, respectively. In contrast, for phenotypes with the interaction term, 1851 and 6076 spuriously significant SNPs were detected for *n* = 100, 000 and *n* = 210, 145 datasets, respectively.

Another limitation of our model is that true *ρ* cannot be obtained without knowing the true interaction architecture. We assessed *ρ*_*max*_ estimates over interaction matrices with higher-order terms and observed that mean *ρ*_*max*_ estimates are consistent but extreme *ρ* values might be less likely with higher-order products (see Appendix B). This is still not sufficient to deduce meaningful conclusions on the range of the *ρ* parameter since the true interaction terms might be defined by a multitude of different nonlinear functions. Without more precise estimates for the range of *ρ*, it is not trivial to transform our model into a diagnostic framework in real GWAS. However, since *ρ* can reach sufficiently high values even when the target SNP and interaction SNPs (from two-way to ten-way, see Appendix B) are on different chromosomes, it is reasonable to assume that real GWAS is affected by this bias. We leave investigating the extent of this bias and developing diagnostic frameworks to future work.

GWAS are crucial for providing the basis of functional studies which are required to understand the causal mechanisms behind complex traits. In this work, we demonstrated how GWAS with linear models can be susceptible to producing spurious associations when the true genetic model has epistasis, especially with increasing sample sizes and when the SNPs involved in epistasis are on the same chromosome. Our findings suggest that future studies should pursue the ongoing efforts of explicit modeling of interactions or assumption-free (or minimum-assumption) models to characterize genotype-phenotype associations [23].

## Contributions

Conceptualisation: BY. Data curation: BY, MNG. Methodology: BY. Investigation: BY, MNG. Formal analysis: BY, GC, FJ. Visualisation: BY. Supervision: BY, MM, GC, FJ. Original draft: BY. Review & editing: BY, TK, FJ, GC.

## Acknowledgements

Thanks to the High Performance Computing Center (HPC) of the University of Tartu for providing computational resources. Thanks to the Estonian Biobank Research Team (Andres Metspalu, Lili Milani, Tõnu Esko, Reedik Mägi, Mari Nelis, Georgi Hudjashov) for the genotype and phenotype datasets used in this work. BY, MNG were supported by the Estonian Research Council grant PSG1179. FJ was supported by the Agence Nationale de la Recherche grant ANR-20-CE45-0010-01. The research was conducted using the Estonian Center of Genomics/Roadmap II funded by the Estonian Research Council (project number TT17). The research was conducted using the research infrastructure “Estonian Center for Genomics” funded by the Estonian Research Council (project number TARISTU24-TK19).

## Ethics statement

The activities of the EstBB are regulated by the Human Genes Research Act, which was adopted in 2000 specifically for the operations of the EstBB. Individual level data analysis was carried out under ethical approval 1.1-12/624 (issued 24.03.2020), 1.1-12/2618 (issued 04.08.2022) and 1.1-12/3454 (issued 20.10.2022) from the Estonian Committee on Bioethics and Human Research (Estonian Ministry of Social Affairs).

## Appendix

### A Variance variance

We prove here that

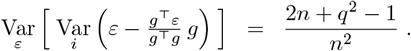

By developing:

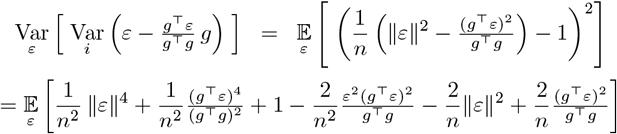

The two last terms were already computed (for the estimation of the average). For the other terms, which are moments of order 4 in *ε*, we will rely on the following formulas for multivariate Gaussian distributions, that is when *X ∼* 𝒩 (0, Σ):

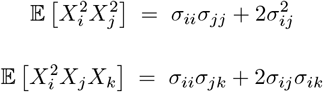

Hence, regarding the first term:

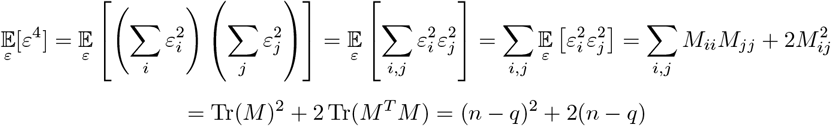

using *M*^*T*^ *M* = *M* .

Regarding the second term, we use the same formula, but using *M* ^*′*^ = *gg*^*T*^ instead of *M* (note that *M* ^*′*^ is symmetric and that *M* ^*′*2^ = ∥*g*∥^2^*M* ^*′*^):

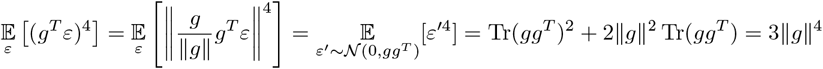

as 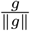 is a unit-norm vector and *g*^*T*^ *ε* is a scalar.

Finally, regarding the third term:

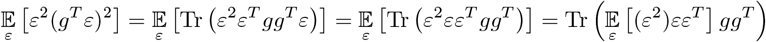

Then 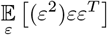 is the following matrix:

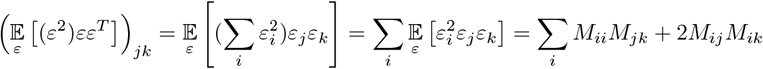

hence

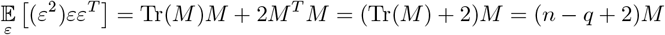

and the third term rewrites:

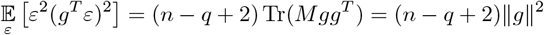

as *Mg* = *g*.

Summing all terms, we have:

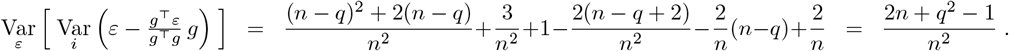

### B Multi-way Interactions

To understand the behavior of *ρ*_*max*_ estimates over higher-order interactions, we obtained *ρ*_*max*_ distributions based on *Z* interaction feature matrices constructed with different N-way interaction terms (i.e., products of centered genotype vectors from N positions), using the same approach as described in Materials and Methods (i.e., *Z* was generated randomly for 100 times with 100 N-way interaction terms, with the constraint that the target SNPs *g* are not part of these features). Regardless of whether the interaction SNPs and target SNPs are on the same chromosome (design 1) or different chromosomes (design 2), mean *ρ*_*max*_ did not change substantially with increasing N. However, variance tended to decrease for design 1 whereas it tended to slightly increase for design 2 with increasing N (Fig 5).

**Figure 5.**
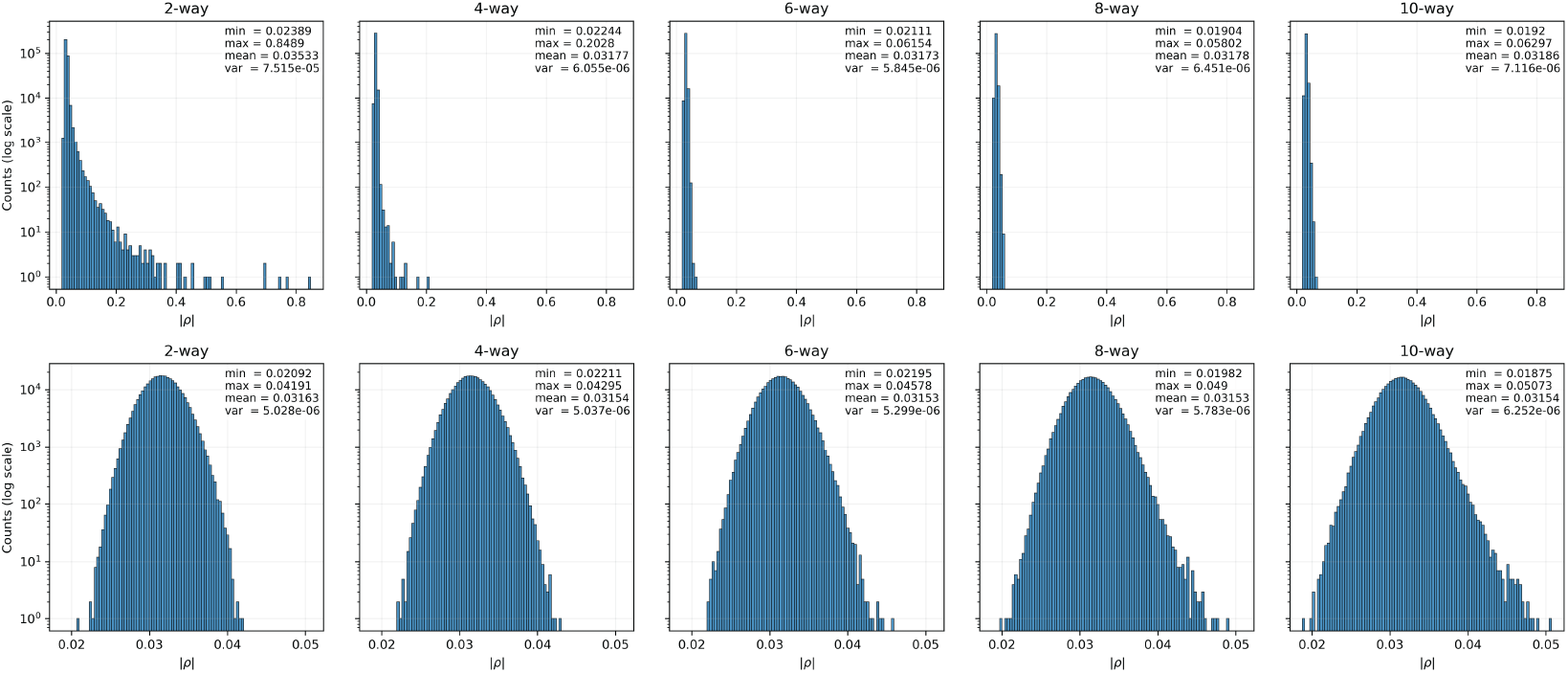
Comparison of the distribution of *ρ*_*max*_ estimates for N-way (N = 2, 4, 6, 8, 10) interaction terms based on designs where the interaction SNPs and target SNPs are on the same chromosome (up) and on different chromosomes (down).

